# Dancing to the beats of the mammalian connectome: Topological and dynamical asymmetries shape infra-slow oscillations

**DOI:** 10.64898/2026.07.01.735524

**Authors:** Anagh Pathak, Rishabh Bapat, Arpan Banerjee

## Abstract

Flexible brain network dynamics unfold on several timescales, despite being constrained by a relatively static structural connectome. Recent cross-species evidence has described infra-slow network modulation, < 0.1 Hz, through two distinct but fundamentally aligned frameworks: transitions between states of integration and segregation identified by high and low global coherence respectively, and periodic transitions between internally and externally oriented attention indexed by the dynamic modulation of alpha power. Here we propose that these infra-slow, structured fluctuations in network engagement stem from a common underlying dynamical motif. Using a network of coupled oscillators embedded in human and macaque empirical connectomes, we show that topological features of brain organization such as modularity, hierarchy and intrinsic dynamical asymmetries naturally give rise to low-frequency collective modes that could nest high frequency states of neuronal communication. These “breathing” dynamics, analogous to beat phenomena in acoustic systems, produce infra-slow fluctuations in global synchrony. Disrupting the connectome topology abolishes such slow coherence oscillations, indicating their dependence on network topology. Furthermore, analytical results from reduced oscillator models reveal how small frequency differences between weakly coupled modules generate slow coherence oscillations, highlighting the importance of dynamical asymmetry. Finally, how neuromodulatory inputs can tune these emergent timescales is discussed, providing a mechanistic link between structural architecture and the dynamic regulation of large-scale brain function.

## Introduction

Large-scale brain networks exhibit remarkable dynamical diversity, communicating across multiple frequency bands to support a wide range of cognitive states^1,2^. Fast network reconfigurations are embedded within slower fluctuations in global engagement, suggesting a hierarchical organization of timescales in cortical dynamics^3,4,5^. This flexibility stands in stark contrast to its structural substrate—the white-matter connectome—which evolves only slowly, over developmental or longer timescales involving months to years. This apparent mismatch raises a fundamental question: how can a relatively stable anatomical architecture give rise to such a rich repertoire of functional dynamics? One promising resolution to this conundrum is that neural dynamics do not simply mirror structure; rather flexible transitions between states of functional coordination are shaped by context^6^ and the modulation of local neuronal interactions^7,8^. Within this view, the connectome constrains a space of possible dynamics, while specific regimes of inter-areal coupling and excitability reveal distinct dynamical motifs embedded within that structure^9,3^. Identifying such motifs, and characterizing the conditions under which they emerge, is therefore key in linking anatomy to function.

The coexistence of functional segregation and functional integration is widely considered a core principle by which brain circuits process information, providing a theoretical framework for understanding complex neuroscientific phenomena such as perceptual binding, sensory integration, conscious awareness, and higher-order cognition^10,11,12,13,14^. Recent studies have identified that the balance between the two is dynamic, realized as distinct temporal periods when the brain exhibits heightened segregation or integration^7,13,15,16^. Increases in integration measures such as global coherence have been associated with improved performance in perceptual binding^17^ or higher-order processes such as working memory and reasoning^15,18,19^. Similarly, desynchronized brain activity biases computation within modules toward independent, localized, and focused processing. These states are referred to as ‘segregative’ and are associated with rapid perceptual and motor functions^14^. Importantly, the balance between integration and segregation is not static, but fluctuates over slow timescales, with prominent spectral components below 0.1 Hz^18,20,21,4,22,23^.

Parallel to the integration–segregation axis, brain activity also fluctuates periodically between externally and internally oriented modes of processing^24^. In externally oriented states, neural activity is shaped primarily by incoming sensory input, supporting perceptual and stimulus-driven computations. In contrast, internally oriented states bias activity toward prior knowledge, facilitating prediction, memory retrieval, and pattern completion^24,25,26^. Converging evidence from fluctuating stimulus detection rates^27,28,29^, self-reported attentional focus^30^, mind-wandering dynamics^31^, and HMM-derived attentional states^29^ suggests that transitions between these modes, much like shifts along the integration–segregation axis, occur over slow timescales (*<* 0.1 Hz). Notably, in sustained attention paradigms this switching exhibits a clear periodic structure. While reported frequencies for this periodicity vary across studies, these dynamics consistently fall within the infra-slow regime, indicating a slow, periodic rebalancing between internally and externally oriented processing. Furthermore, such slow periodic switching has been found to be conserved across humans and macaques, surprisingly at similar frequencies^27,28^.

We propose that the commonly invoked dichotomies of integration versus segregation and internal versus external processing may not be independent, but instead reflect complementary descriptions of a single underlying dynamical motif. In this view, integration and segregation capture the computational organization of large-scale neural interactions, whereas internally and externally oriented states represent their behavioral manifestation. This perspective naturally leads to a central question: what is the simplest, most parsimonious dynamical mechanism capable of generating such structured, slow fluctuations between functional states? That is, following David Marr^32^, how can this computation be realized at the level of its dynamical implementation? To address this, we model the brain as a network of coupled Kuramoto oscillators^33^ interacting through the empirical white-matter connectome. Within this framework, we show that the topology of the mammalian connectome intrinsically supports the emergence of slow collective modes. In particular, we identify *asymmetry, modularity*, and *hierarchical organization* as key topological principles that give rise to low-frequency “breathing” dynamics, analogous to beat generation in acoustics (Figure 1a). These dynamics naturally produce slow fluctuations in global coherence, effectively reconciling integration/segregation and internal/external processing modes (Figure 1b,c). Consistent with this interpretation, we further demonstrate that disrupting the connectome through topological randomization abolishes these slow modes, showing that they are not generic features of coupled systems but instead depend critically on structured connectivity. Both the human and macaque connectomes share these topological principles, facilitating the production of this dynamic motif. We further substantiate this mechanism analytically using reduced networks of Kuramoto oscillators, showing how small frequency mismatches between weakly coupled modules generate slow fluctuations in global synchrony. Finally, leveraging this formalism, we demonstrate how neuromodulatory inputs can systematically tune these emergent timescales by interacting with the underlying topology, thereby providing a mechanistic link between structural organization and the dynamic regulation of large-scale brain activity.

**Figure 1:**
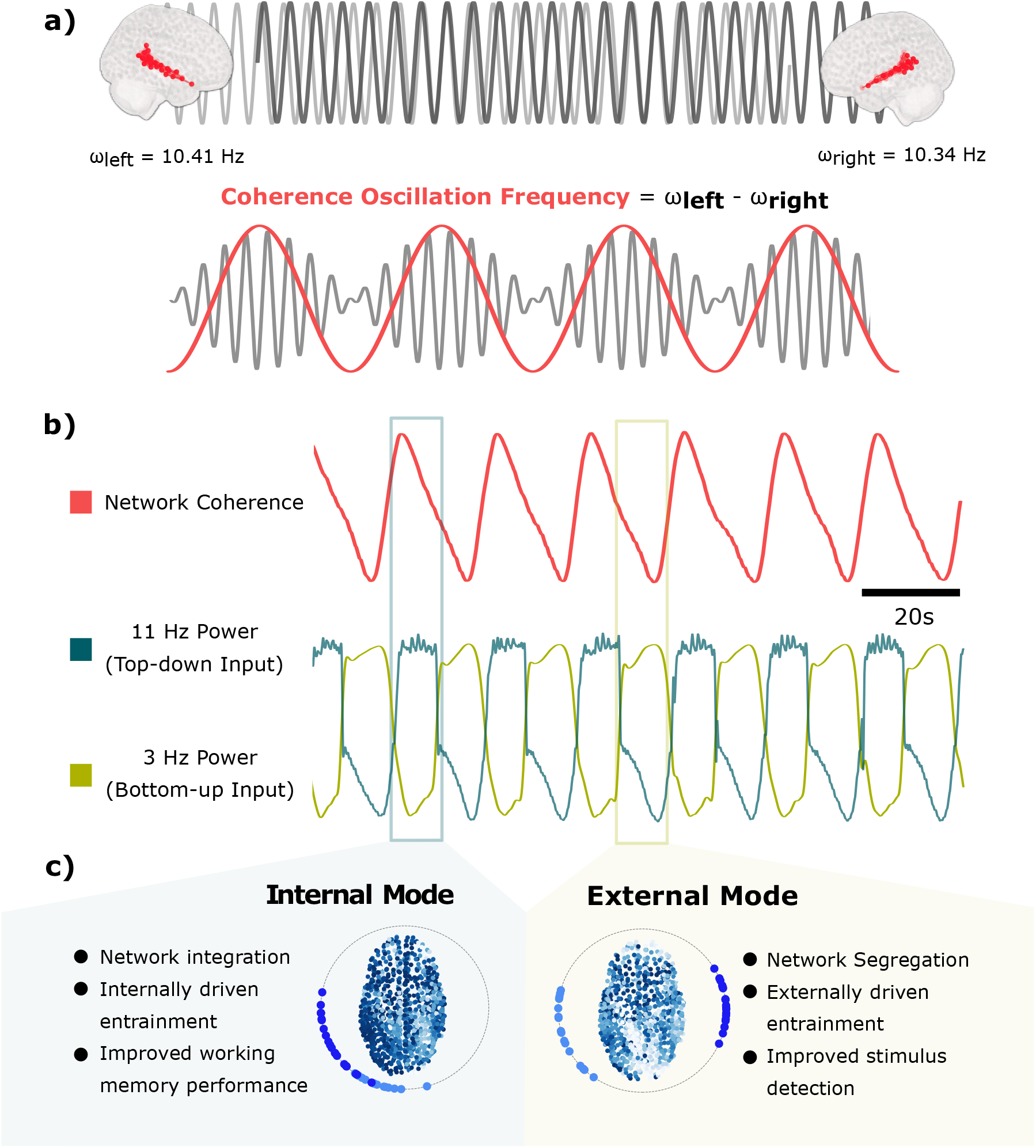
Periodic Integration and internal/external modes: **a)** The functional networks of the brain can be divided into hemispheric modules; here, the temporo-parietal network is shown in red. Each module synchronizes at a slightly different frequency. **b)** Frustrated synchronization between modules leads to periodic integration in the whole network. The current article proposes that top-down input and sensory information compete to entrain the sensory cortex. High coherence increases entrainment by top-down input. **c)** High and low coherence periods could map onto internal and external processing modes, reflecting varying degrees of top-down input onto sensory regions and allowing the brain to optimally use the dual efficiencies of both integrative and segregative processing.

## Results

In order to establish the relevance of our integration/segregation dynamical motif, we begin by situating it within prior empirical findings on rhythmic attentional fluctuations. In a series of studies, Lakatos et al. ^27^ demonstrated that selective attention is not continuous but instead fluctuates rhythmically at very slow frequencies (≈ 0.06 Hz in macaques). Crucially, lapses in externally oriented attention were found to coincide with periods of elevated alpha power and diminished neural entrainment to sensory stimuli. More recent work in humans using EEG has corroborated these findings, revealing remarkably similar slow fluctuations (≈ 0.07 Hz), thereby suggesting a conserved dynamical process across species^28^. Throughout this article, we refer to this dynamical pattern—characterized by the slow alternation between externally and internally oriented processing states—as the *slow coherence oscillation* (SCO) motif. We further hypothesize that such dynamical motifs constitute a fundamental organizational feature of large-scale brain activity and arise from the underlying structural architecture of the brain. Specifically, we model large-scale neural dynamics as emerging from the interaction of coupled oscillatory units, with natural frequencies in the alpha range to remain consistent with Lakatos et al. ^27^. The interaction terms in the coupled oscillator model are derived from metrics of white-matter connectivity estimated from diffusion-weighted imaging used in previous studies^34^. Within this framework, often referred to as the ‘Virtual Brain’ ^34,35,36,37^, the balance between integration and segregation can be mathematically operationalized using the *order parameter*, a concept borrowed from the field of synergetics^38^, with higher values reflecting increased phase clustering (integration) and lower values reflecting reduced phase clustering (segregation). Hence, the emergence of SCOs becomes equivalent to having solutions in which order-parameter evolution is periodic. In the following sections, we investigate whether the order parameter arising from such large-scale dynamics can exhibit oscillatory behavior, and we characterize the conditions and topological constraints under which such solutions emerge.

### SCO motifs emerge naturally in the virtual brain

Under our model, slow periodic changes in cortical integration underlie internal and external processing modes. To test whether periodic changes in integration could emerge from the structural organization of the brain, we construct a model of cortical synchronization dynamics (Figure 2a). The human cortex is simulated as a set of 1000 functionally distinct regions based on the Schaefer^39^ atlas. Each region is modeled as a Kuramoto phase oscillator^33^ with an intrinsic frequency and delayed sinusoidal coupling to other regions (see Simulating Synchronization Dynamics). Inter-regional coupling is scaled based on a structural connectivity matrix derived from empirical tractography, and intrinsic frequencies are distributed from the alpha band (8–12 Hz) based on the core–periphery axis (see Deriving Structural Connectivity Matrices and Assigning Intrinsic Frequencies). Phase synchronization is measured using the Kuramoto order parameter^33^ and used to index integration at the whole-cortex level and in functional networks defined by Yeo et al. ^40^.

**Figure 2:**
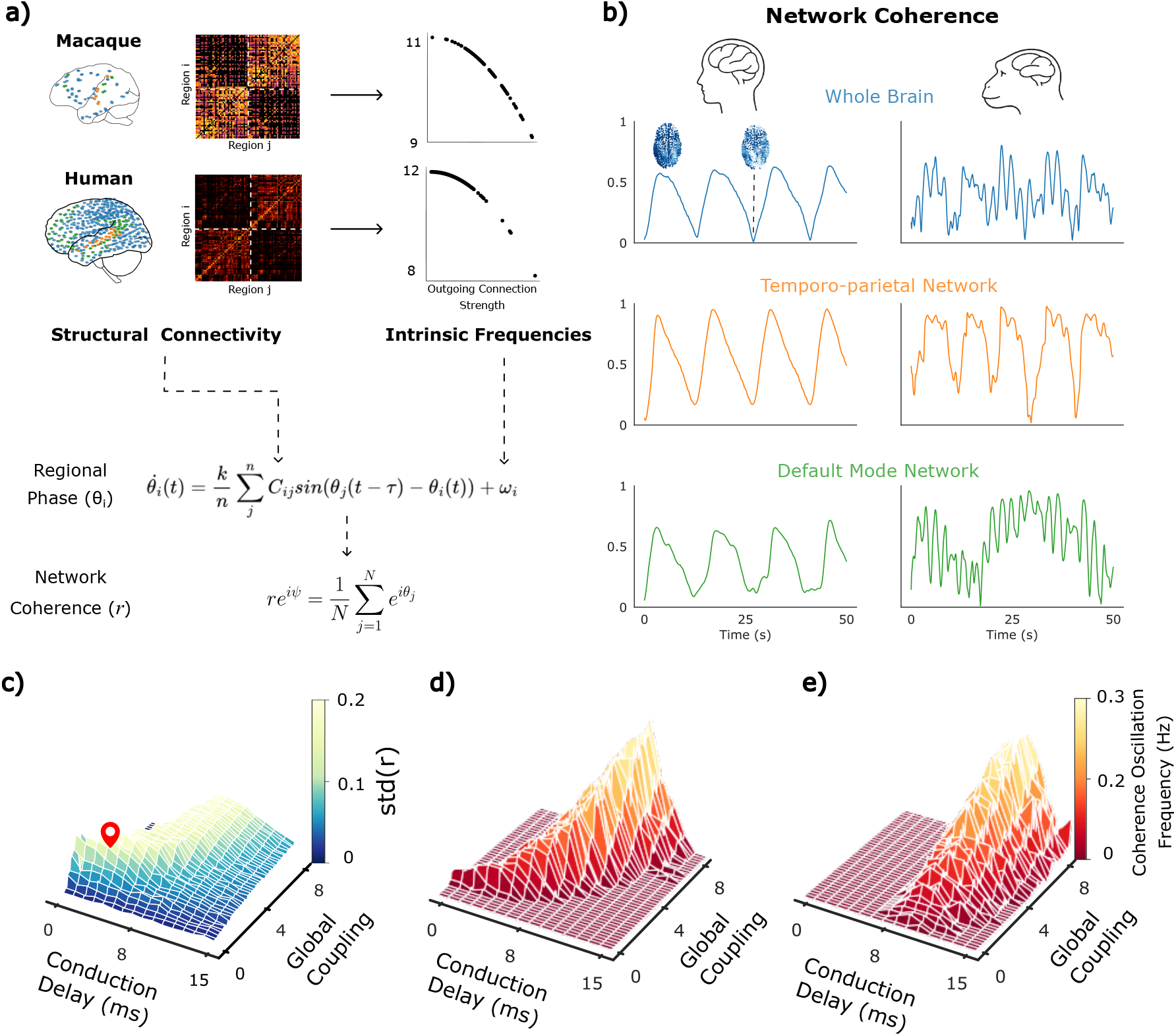
Emergence of periodic integration in the human and macaque connectomes: **a)** An overview of the connectome-based cortical model. A network of Kuramoto oscillators is constructed based on the Schaefer human parcellation^39^ and the regional map macaque parcellation^42^. The coupling between each pair of oscillators is based on empirical tractography and the intrinsic frequencies are drawn from the alpha range based on the strength of outgoing connections from each region. The Kuramoto order parameter measures coherence and is used as an index of network integration. **b)** Coherence time series for the whole cortex, temporo-parietal and default-mode networks. Coherence oscillations are observed in both the human and macaque connectomes. A phase plot where each node is colored based on its phase is provided to illustrate the difference between integrated and segregated states. **c)** Surface plot showing how the standard deviation of cortical coherence varies across the parameter space for the human connectome. This indexes metastability, a feature of resting-state neural activity^46^. Delay and coupling were varied to maximize metastability for both the human (coupling = 2, delay = 4) and macaque connectome (coupling = 2.6, delay = 5). **d)** Surface plot showing how coherence oscillation frequency changes over the parameter space for the human connectome. **e)** The effect of intrinsic frequency heterogeneity on coherence oscillations across the parameter space. For homogeneous intrinsic frequencies coherence oscillations cannot exist at low delays.

The degree of synchronization in the cortical model is influenced by a global scalar for coupling and the amount of conduction delay. The parameter space spanned by these values reveals a region where the model exhibits partial synchronization. In this dynamical regime, the cortical model shows periodic integration globally and in the temporo-parietal and default-mode networks (Figure 2b). To test whether the dynamics are robust to minor structural changes and coarser parcellation, we simulate activity using alternative connectomes. A human connectivity matrix based on the 90-region Automated Anatomical Labeling Atlas^41^ also shows periodic changes in integration over a wide range of parameters. Intriguingly, an 82-region macaque connectome based on the regional map parcellation^42^ exhibits similar dynamics within the whole cortex and in analogs of human functional networks^43,44,45^. This suggests that an evolutionarily conserved aspect of structure may form the basis for periodic changes in integration.

The peak frequency of global integration varies considerably over the parameter space defined by global coupling and delay. To identify the set of parameters most comparable to resting-state neural activity, we maximize the variance in global synchronization by performing a grid search over a range of global coupling and delay values. This is thought to index metastability, a feature of resting-state cortical activity that enables the flexible coordination of neural ensembles^46^. Studies show that maximizing metastability in computational models increases the similarity between simulated and empirical neural dynamics^47^. Our model shows a similar result: the values of global coupling and delay that maximize metastability also produce periodic changes in global integration at ≈ 0.07 Hz in the human connectome, matching empirically observed switching frequencies between internal and external modes.

To further establish the unique propensity of the human connectome to generate oscillations in the order parameter, synchronization dynamics were simulated in two alternative synthetic networks (see Methods),

1. A fully connected network with heterogeneous natural frequencies drawn from a Lorentzian distribution (*n* = 1000)
2. Randomized human (*n* = 1000) and macaque (*n* = 82) connectomes, with intrinsic frequencies drawn from the alpha range (see Assigning Intrinsic Frequencies).

Notably, for scenario 1, analytical solutions exist for describing the evolution of the global order parameter. Specifically, for *N* → ∞ it has been shown^34,48^ that the order parameter for a frequency distribution given by,

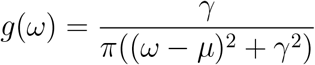

reduces to the following in the absence of delays,

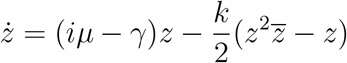

where *z* is the complex order parameter. To obtain the real-valued order parameter, substitute

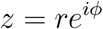

giving,

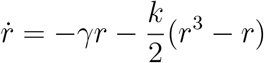

As a one-dimensional ordinary differential equation, this system does not admit oscillatory solutions. This is confirmed through numerical simulation with *n* = 1000 oscillators. Simulations further preclude order-parameter oscillations for idealized all-to-all networks with homogeneous conduction delays (Figure 3b). Numerical simulations further indicate that scrambled human or macaque connectomes, formed by randomly shuffling connection weights from the original connectome while preserving bi-directional symmetry, do not yield oscillations in the global order parameter (Figure 3a).

**Figure 3:**
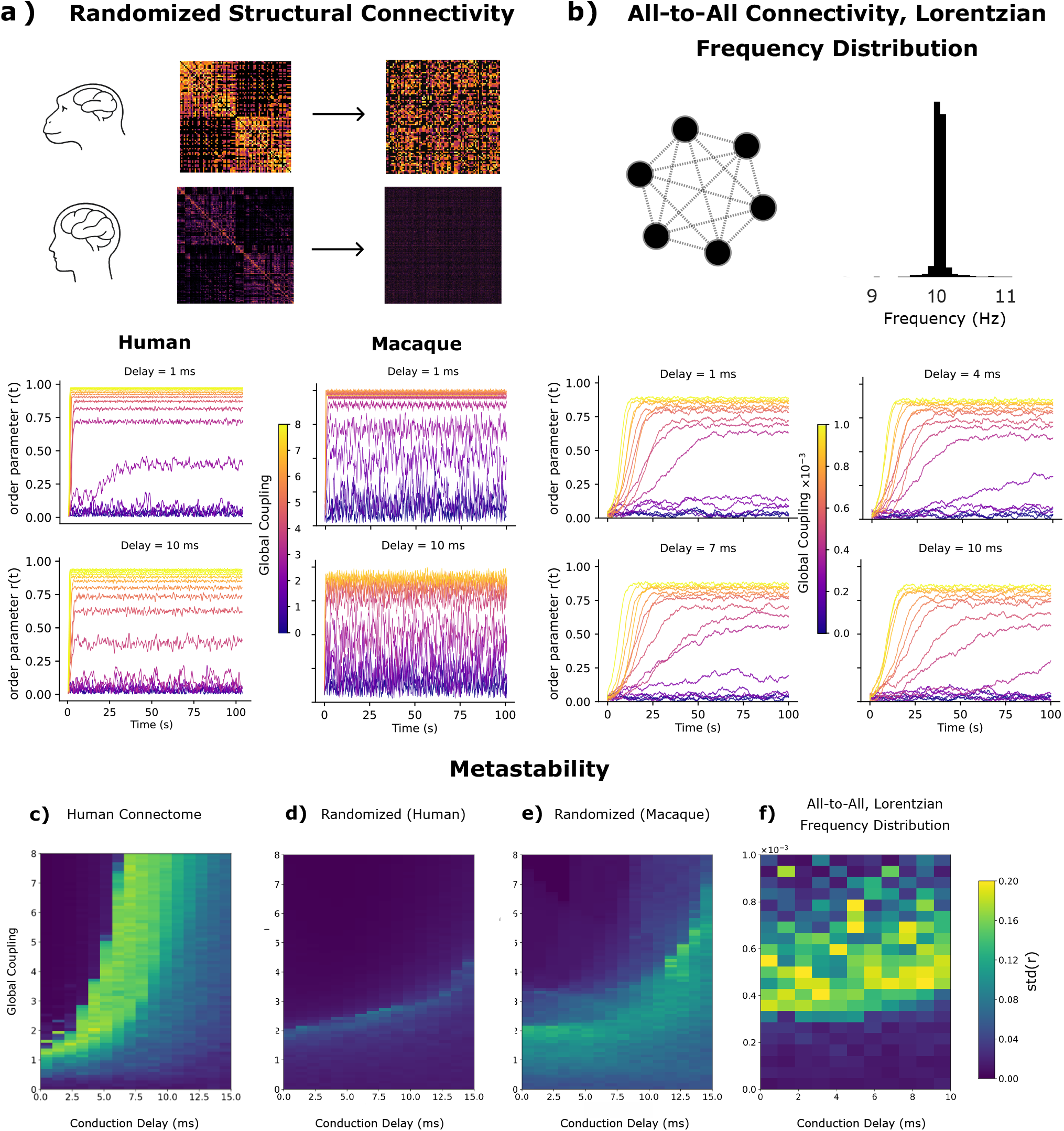
Order parameter for null models: **a)** Human and macaque connectomes were randomly permuted while retaining connection strengths. The model was otherwise unchanged from previous analysis. **b)** An all-to-all network with 1000 Kuramoto oscillators and natural frequencies following a Lorentzian distribution. **(Below)** Corresponding order parameter time series for both networks at various values of global coupling (represented by color) and at various (homogeneous) delays. The resulting order parameter time series show no periodic structure. **c), d), e), f)** Metastability over the parameter space spanned by global coupling and delay. Randomization reduces metastability across the parameter space for human and macaque connectomes. An all-to-all connectome shows a similar range of metastability values to human and macaque connectomes; however, the order parameter dynamics lack a periodic structure.

### Dynamical basis of periodic integration

Next, we seek to understand the underlying principles that give rise to SCOs. We begin by considering a simple network consisting of two populations of Kuramoto^33^ oscillators (Figure 4a). The network has a modular connection topology, with strong intra-modular coupling values (*k*_*a*_ and *k*_*b*_) and weak inter-modular connectivity (assumed to be zero for the sake of simplicity). The oscillators within each module also have intrinsic frequencies (*w*_*a*_ and *w*_*b*_), and connections are subject to delays (*τ*_*a*_ and *τ*_*b*_). Due to the modular connection topology and an appropriate strength of connectivity, the two populations synchronize locally (*r*_*a*_ = *r*_*b*_ = 1), but not globally (*< R >* ≈ 0), in a phenomenon known as frustrated synchronization^49^. Furthermore, if the modules synchronize at different frequencies (for instance, due to varied distributions of natural frequencies and/or delays), the global order parameter exhibits oscillations. The frequency of this oscillation is equal to the difference between the synchronization frequencies of the two modules. This idea is directly analogous to *acoustic beats*, where the interference between two nearby frequencies generates an envelope at their difference frequency.

**Figure 4:**
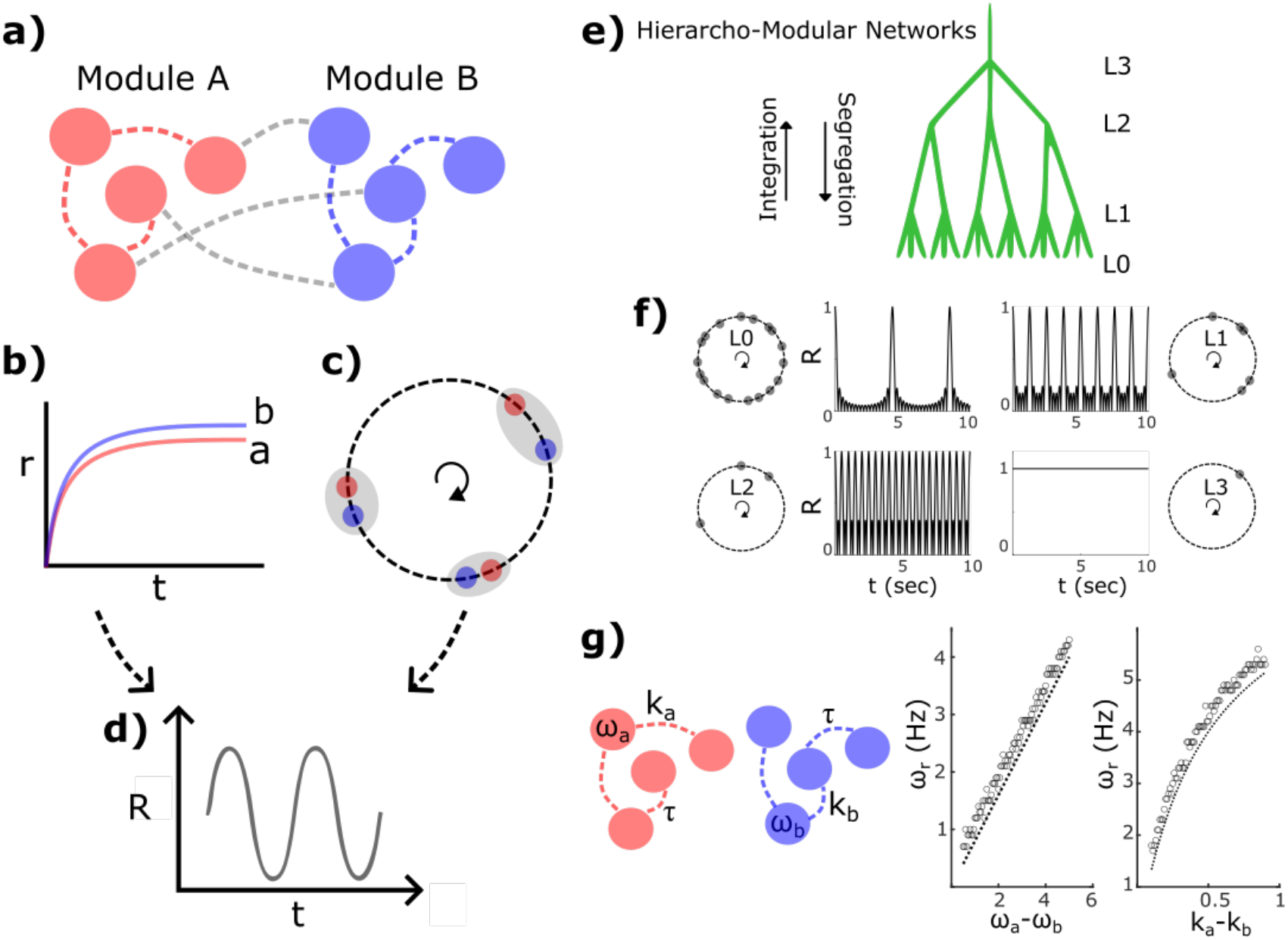
Frustrated synchronization and periodic order parameter: The emergence of oscillating order parameters may be understood through a simplified toy model: **a)** two modules with strong within-network coupling and relatively weak between-network coupling; **b)** complete within-network but no between-network synchronization; **c)** two modules with different internal synchronization frequencies; and **d)** an oscillating order parameter estimated over the entire population. This phenomenon may be generalized to any hierarcho-modular (HM) network. For example, **e)** an HM network consisting of 27 oscillators with uniformly distributed natural frequencies between 8–12 Hz; **f)** at level *L*_0_, the network is completely asynchronous, *L*_1_ corresponds to internal synchronization of sub-modules in groups of 3, and similarly for higher levels *L*_2_ and *L*_3_; **g)** frequency dispersion and delays can influence modular synchronization frequency. **(right)** Order-parameter frequency as a function of frequency dispersion or coupling; dotted lines represent analytical expressions, while circles represent numerical results with 500 oscillators in each module.

Under suitable approximations, the synchronization frequency for each module can be derived analytically, offering further insight into the mechanism underlying this beat phenomenon. Specifically, consider a network of *n* identical Kuramoto oscillators (natural frequency = *ω*_0_, delay = *τ*). Each oscillator is governed by the following,

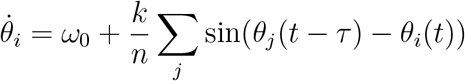

For strong coupling, the network synchronizes completely such that each node settles at a common frequency *ω*_*s*_. In the steady state, the phase of each node evolves as,

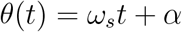

Substitution yields,

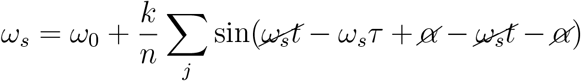

For small delays,

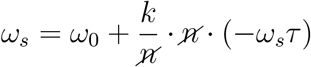

This yields an expression for the synchronization frequency,

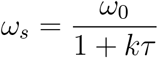

Consequently, the frequency of oscillation in the global order parameter (Figure 4d) is given by the following expression.

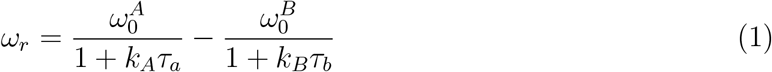

Inspection of the above expressions reveals the key mechanisms governing the resulting beat frequency. In this simplified framework, differences in intrinsic frequencies, coupling strengths, and delays introduce asymmetries between the two modules, which in turn determine the magnitude of the beat frequency, either amplifying or attenuating it. This perspective helps explain why SCOs emerge over a broader range in the human connectome when heterogeneities in natural frequencies are introduced. Such heterogeneity increases dynamical frustration, thereby enhancing frequency detuning between modules (compare Figure 2d with Figure 2e). Notably, these asymmetries are not artificially imposed but are inherent to mammalian connectomes. Assigning natural frequencies as a function of degree further accentuates these effects by systematically introducing additional asymmetry. Importantly, in the absence of delays (*τ*_*a*_ = *τ*_*b*_ = 0), the denominator reduces to unity, effectively diminishing the influence of coupling asymmetries. This highlights that delays, coupling, and intrinsic frequency differences act synergistically to shape the resulting dynamics. We leverage this insight in a later section to demonstrate how modulation of network connectivity can tune dynamical asymmetries. Building on this insight, the same mechanism extends naturally to hierarchical systems: when modules are organized across multiple levels, frequency detuning at each level gives rise to a cascade of beat interactions, generating multiple emergent timescales. Consequently, the temporal structure of the system reflects its hierarchical organization. We demonstrate this principle directly in Figure 4e,f.

Extending this to the mammalian brain, its hierarchical organization implies that the same mechanism operates recursively across scales: sub-networks embedded within larger networks exhibit similar dynamics in a self-similar manner. This property is recapitulated in our cortical model, where both the full cortex and its constituent sub-networks display oscillatory behavior in their respective order parameters (Figure 2b). Within each sub-network, locally synchronized modules are organized along hemispheric lines. For instance, in the temporo-parietal network, the left and right sub-networks may synchronize at slightly different frequencies. Their interaction then gives rise to a slow oscillation in the global order parameter, with a frequency equal to the difference between the synchronization frequencies of the two hemispheric modules—directly analogous to a beat phenomenon in coupled oscillatory systems. More generally, such partially synchronized, frustrated states are consistent with the notion of a Griffiths phase, characterized by an extended regime of critical-like dynamics.

### Competitive interactions between network dynamics and external stimuli

Lakatos and colleagues^27^ demonstrated that sustained attention is not continuous but fluctuates rhythmically at infra-slow frequencies (≈ 0.06 Hz), with alternating periods of strong sensory entrainment and disengagement. Crucially, these lapses of externally oriented processing are accompanied by elevated alpha power, consistent with a shift toward internally oriented or inhibitory states. More recent work in humans has extended these findings, showing that opposing processing modes—characterized by high entrainment to external stimuli versus increased alpha activity—alternate rhythmically over time and are tightly coupled to behavioral performance^28^. Together, these studies establish that attention is governed by structured, slow fluctuations in which alpha and entrainment exhibit an antagonistic relationship, reflecting shifts between internal and external modes of processing. While the phenomenology of these dynamics is now well documented, their mechanistic origin remains unclear. We suggest that such alternations may arise naturally from interactions between weakly detuned network modules, which generate slow coherence oscillations through a beat-like mechanism. In this view, periods of high global coherence correspond to internally dominated states marked by elevated alpha, whereas periods of reduced coherence favor external entrainment. Thus, SCOs provide a dynamical systems account linking large-scale network organization to the rhythmic interplay between alpha activity and sensory processing observed during sustained attention.

To account for this interplay, we consider a cortical region receiving both feedforward sensory input and feedback from an ipsilateral sensory network (e.g., the temporo-parietal network in the auditory domain). Within this framework, external sensory drive and ongoing network activity compete for representation within the sensory region, giving rise to two distinct dynamical regimes (Figure 5a). During periods of high temporo-parietal coherence, the network exerts a unified influence on the sensory node, effectively entraining it to the intrinsic network rhythm. These epochs are associated with elevated alpha power both locally and across the broader network, consistent with reports of increased alpha coherence during attentional lapses^27^. In contrast, during periods of low temporo-parietal coherence, the opposing influences of left and right sub-networks partially cancel, reducing net feedback and allowing external input to dominate (Figure 5b). These periods are characterized by enhanced entrainment to the sensory stimulus, reflected in increased power at the driving frequency. Importantly, transitions between these internal and external processing modes unfold at the infra-slow timescale set by the SCO (≈ 0.07 Hz). Furthermore, the in-phase coupling between local temporo-parietal coherence and whole-brain integration (Figure 2b) gives rise to dynamic routing states, providing a systems-level mechanism for dual-mode attentional processing.

**Figure 5:**
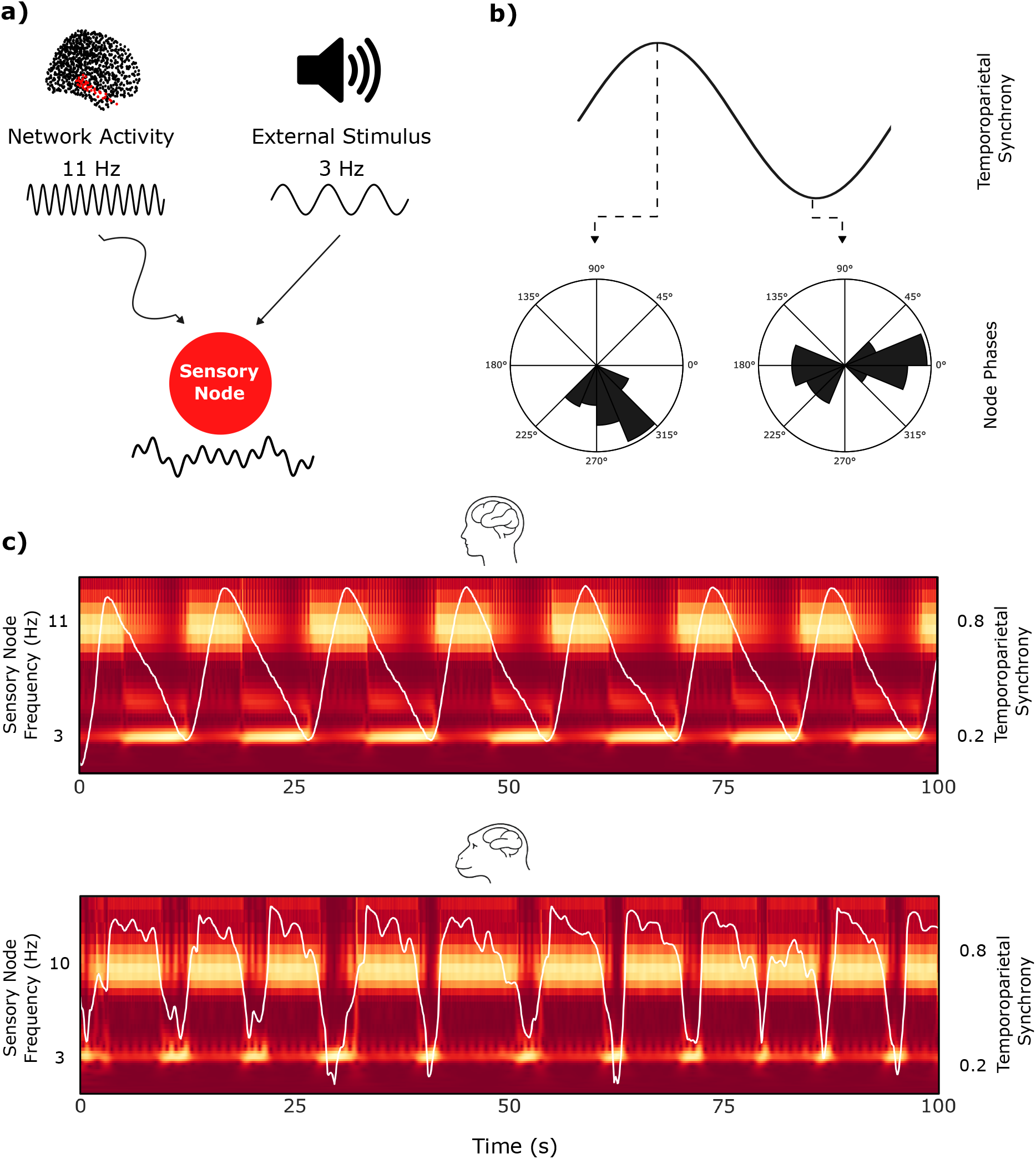
Competitive entrainment in the cortical model: **a)** Schematic illustrating the competitive entrainment model used to relate oscillations in coherence to internal and external processing modes. A simulated sensory node is coupled to the entire temporo-parietal network (whose activity is at 11 Hz) and to a 3 Hz stimulus. **b)** Polar histograms showing the node phases of the temporo-parietal network during periods of high and low coherence. When synchrony is high, the temporo-parietal network exhibits tightly clustered node phases and thus exerts a stronger influence on the sensory node. **c)** Time-frequency plot of the simulated sensory node (brighter means more power at the frequency on the left y-axis). The white line and the right y-axis show the temporo-parietal coherence over time. The node oscillates in the alpha band when temporo-parietal coherence is high and at the stimulus frequency when temporo-parietal coherence is low. Similar results exist for a cortical model based on the macaque connectome.

### Tuning the Timescale of Periodic Integration

We next consider whether the frequency of SCOs could be dynamically modulated as a function of task demands. As established above, SCOs arise from subtle structural asymmetries between network modules, which shape their respective synchronization frequencies. Any mechanism capable of modulating the effective expression of these asymmetries should therefore be able to tune SCO frequency. One straightforward route is to alter the degree of functional asymmetry between modules, thereby shifting their synchronization frequencies and, consequently, the resulting beat frequency that underlies the SCO. A growing body of work demonstrates that neuromodulatory systems, particularly cholinergic and noradrenergic projections, can selectively tune the balance between functional integration and segregation through regionally specific modulation of neural gain and coupling^7,16^. Crucially, these effects are not spatially uniform but depend on receptor density and network embedding, enabling the selective amplification or suppression of functional asymmetries across modules.

Within this framework, increased cholinergic tone could amplify lateral asymmetries in the temporo-parietal network (Figure 6a), thereby increasing the separation between module-specific synchronization frequencies and modulating the timescale of periodic integration (Figure 6b). To test this mechanism, both human and macaque cortical models were initialized near maximal metastability (Figure 6c–f). Coupling strength was then selectively increased in either the left or right temporo-parietal sub-network (see Simulating Cholinergic Gain), effectively enhancing lateral asymmetry (Figure 6a). From Equation 1, increasing a module’s coupling (*k*_*A*_ or *k*_*B*_) while holding delays fixed lowers that module’s synchronization frequency. Applying gain to the faster-synchronizing (left) sub-network therefore reduces the detuning between modules and slows the SCO (Figure 6b,e), whereas applying gain to the slower (right) sub-network increases the detuning and speeds it up (Figure 6d,g). However, the system requires conduction delays (*τ* ≠ 0) for gain changes to modulate the SCO frequency.

**Figure 6:**
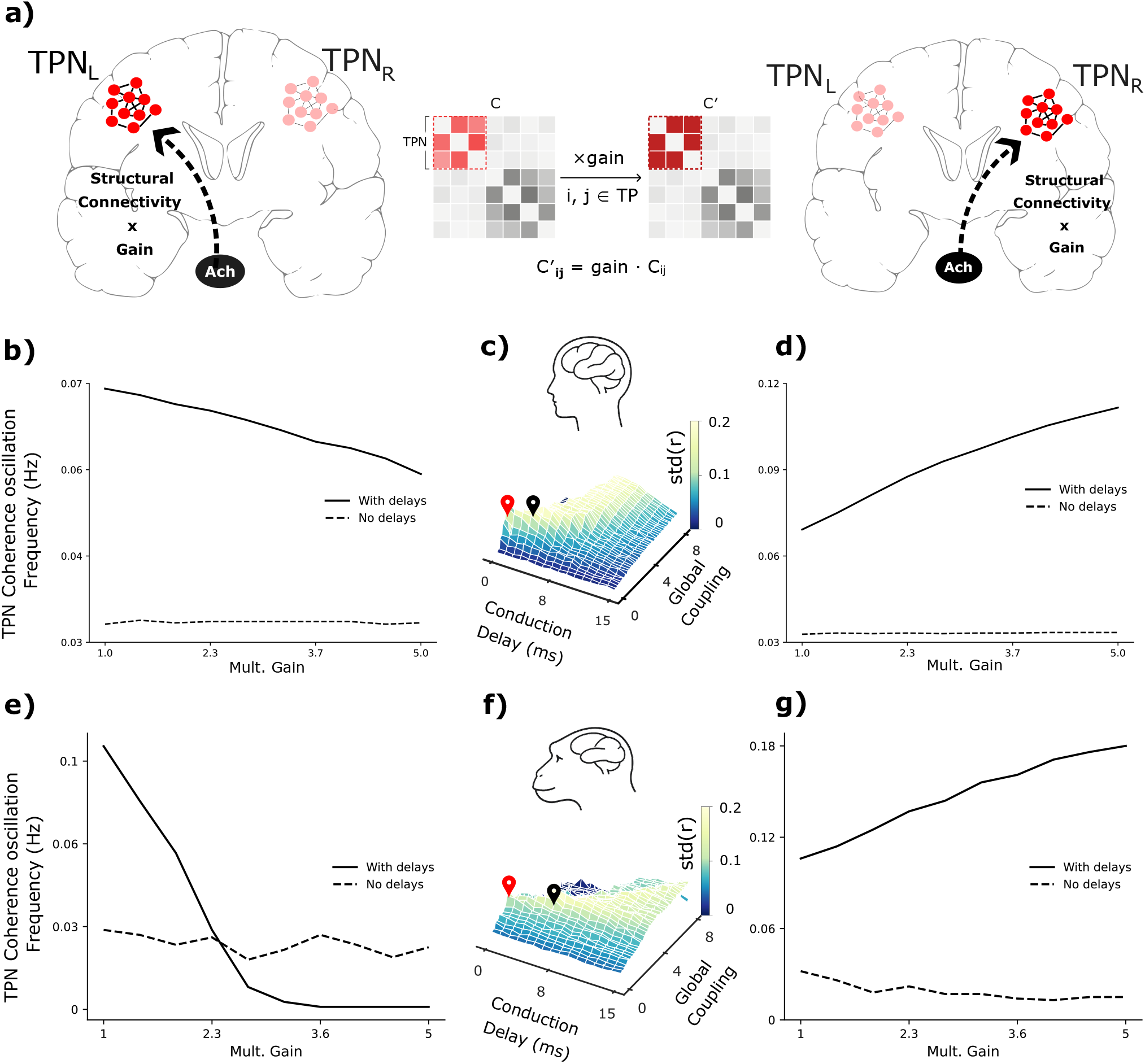
Cholinergic modulation: **a)** Schematic illustrating multiplicative gain applied to the left and right hemispheric temporo-parietal networks (TPN). The gain is multiplied with the structural connectivity matrix at indices within each sub-network. From Equation 1 this is expected to lower the synchronization frequency of the stimulated sub-network. **b), d)** The effect of gain applied to the left and right temporo-parietal networks for the human connectome. Since the left sub-network synchronizes at a higher frequency than the right sub-network, when gain is applied to the left sub-network the difference in their synchronization frequencies reduces and the temporo-parietal coherence oscillation slows down. The opposite effect occurs when gain is applied to the right temporo-parietal network. As predicted by Equation 1, the absence of delays makes it impossible for cholinergic gain to affect periodic integration frequency. **e), g)** Similar results exist for the macaque connectome. **c), f)** The points in the parameter space used for simulation in the human and macaque connectomes. The delayed condition is indicated by a black marker while the non-delayed condition corresponds to the red marker. The color bar indicates the standard deviation of the order parameter (*r*), which is a measure of metastability^46^.

## Discussion

The key goal of the current study is to demonstrate that the mammalian connectome is inherently predisposed to support dynamical regimes wherein states of global synchrony fluctuate at infra-slow timescales. Through a combination of null modeling, analytical insights, and mechanistic reasoning, we identify the topological and dynamical features underlying this phenomenon, which we term slow coherence oscillations (SCOs). In particular, the connectome is hierarchically modular and subtly asymmetric at the level of sub-modules. Within a partially synchronized regime, emerging at intermediate levels of coupling, these sub-modules synchronize at slightly different frequencies. The resulting frequency detuning gives rise to slow fluctuations in global coherence, a mechanism that is naturally understood in terms of an acoustic beat analogy, where the superposition of nearby frequencies produces slow amplitude modulations. By studying limiting cases in which inter-modular structure is removed, we also show that modular architecture and non-zero time delays are necessary ingredients for the emergence of SCOs. Finally, we show that cholinergic gain modulation can tune SCO frequency by introducing asymmetric coupling among two or more modules.

A mechanistic theory of SCOs provides a parsimonious explanation for cyclical alternations between integration and segregation reported by previous studies^18^. Periods of high global coherence correspond to states in which network activity exerts a strong, unified influence over local regions, favoring internally oriented processing^28^. In contrast, periods of low coherence reflect a relative decoupling of local nodes from large-scale dynamics, increasing their susceptibility to external input and thus promoting externally oriented processing modes. In this way, SCOs offer a dynamical substrate for rhythmic switching between internal and external processing states. The idea of network motifs is not new and has been used to provide phenomenological explanations of biological processes^50^. Borrowing this conceptual inspiration, we go beyond a heuristic description of the phenomenon by using well-established models of whole-brain neuronal dynamics, such as the Kuramoto system.

An important outcome of our analysis is the demonstration that a stable structure need not make the timescales of functional segregation/integration or external/internal modes of processing fixed; rather, they can be flexibly tuned. Specifically, neuromodulatory systems may regulate SCO frequency by modulating the effective asymmetry between network modules, thereby altering their synchronization frequencies and the resulting coherence dynamics. This provides a plausible mechanism through which the brain can adaptively control the temporal structure of large-scale coordination in response to changing task demands. Notably, our analysis further indicates that this modulatory mechanism critically depends on the presence of finite conduction delays, highlighting that such delays are not merely a biophysical constraint but a functional ingredient of large-scale brain dynamics. In this sense, delays play an active role in shaping the temporal structure of coherence fluctuations and enabling the emergence of tunable SCOs. More broadly, this observation aligns with longstanding views that the brain leverages its underlying geometry, topology, and spatiotemporal embedding to support computation at the systems level^9,51,52^. Internally generated attentional switching, recently demonstrated behaviorally by Glukhova et al. ^53^ across three species, provides direct predictive validation of the framework presented here. In their results, attentional states switch rhythmically, with conserved dwell times and occupancy patterns across species. Transitions between attentional states occur in the absence of any externally imposed rhythm and show no systematic relationship to task difficulty. This is consistent with the view that periodic integration, and the attentional states it relates to, are emergent properties of connectome topology.

A prominent axis of asymmetry in the human connectome is hemispheric laterality. Beyond global left–right differences, many large-scale functional systems—including the temporoparietal network (TPN) and default-mode network (DMN)—exhibit pronounced lateralized organization^54,55,56,57,58^. Such lateralization is a conserved feature across species and has long been associated with cognitive advantages, including increased processing efficiency, reduced redundancy, and the capacity for functional specialization (e.g., language lateralization, spatial attention biases, and motor dominance). These observations have led to the prevailing view that hemispheric asymmetry serves primarily to partition cognitive functions across distributed neural substrates. Our results suggest a complementary and more dynamical role for lateralization. In our model, asymmetries between homologous sub-networks give rise to slight differences in their synchronization frequencies, which in turn generate slow coherence oscillations (SCOs) through a beat-like mechanism. From this perspective, hemispheric asymmetry does not simply partition function across space, but also endows the system with an additional *temporal dimension*, enabling the emergence of slow, structured fluctuations in large-scale coordination. Thus, lateralization may serve as a fundamental computational mechanism, giving rise to tunable infra-slow dynamics that regulate the balance between integration and segregation.

This interpretation has direct implications for attention. As shown here, SCOs provide a dynamical substrate for rhythmic alternations between internally and externally oriented processing states. Given that hemispheric asymmetry is a key driver of these oscillations, disruptions to lateralized organization would be expected to impact the stability or frequency of these dynamics. We therefore speculate that atypical hemispheric asymmetry may contribute to attentional impairments by altering the temporal structure of large-scale coordination. Consistent with this view, recent work has reported altered indices of structural and functional asymmetry in individuals with Attention Deficit Hyperactivity Disorder (ADHD), including differences in cortical thickness, white-matter organization, and inter-hemispheric connectivity^59,60^. While these findings are typically interpreted in terms of impaired specialization or inefficient communication, our framework suggests an additional possibility: that such alterations may disrupt the generation or tuning of SCOs, thereby impairing the rhythmic coordination of internal and external processing modes that underlies sustained attention.

## Conclusions

Periodic alternation between internal and external attentional modes is a prevalent theme in the literature that lacks a comprehensive explanation. We propose that switching between integrative and segregative states is the dynamical principle behind this phenomenon, and that competitive entrainment is the mechanism relating them. Computational models demonstrate that both human and macaque connectome-based whole-brain models can support periodic integration at frequencies relevant for alternation of internal and external modes of attentional processing. Further simulations show that, for a given sensory region, periodic changes in stimulus entrainment could emerge from competition between the sensory network and an external stimulus. Modularity is revealed as the structural basis for periodic integration, and conduction delays are adaptive because they enable cholinergic gain to modulate periodic integration frequency. Some important factors remain outside the scope of our discussion. In particular, the roles of cross-frequency coupling and thalamo-cortical interactions in sensory gating are not explicitly modeled. Nonetheless, our results bring a wide variety of findings under the same theoretical framework and suggest exciting new directions for the computational modeling of adaptive cognition.

## Methods

### Deriving Structural Connectivity Matrices

Weighted structural connectivity matrices were derived from empirical tractography in humans and macaques. The human connectivity matrix was obtained from a public repository maintained by Deco et al. ^47^. Diffusion spectrum MRI data of 32 young adults from the human connectome project^61^ were used to determine fiber orientation distributions using q-sampling^62^. The fiber orientation distributions were registered to white matter masks derived from T2-weighted MRI, and deterministic tractography^63^ was used to construct anatomically constrained tractograms. The tractograms were then transformed to Montreal Neurological Institute space^64^ and mapped onto the 1000 cortical regions defined by the Schaefer parcellation^39^. 1000 by 1000 connectivity matrices were then derived for each subject based on the fiber density between each pair of regions. The subject-averaged connectivity matrix was used to fit an exponential distance rule to arrive at the final structural connectivity matrix^47^.

The macaque structural connectivity matrix was uploaded publicly by Shen et al. ^42^. Diffusion-weighted MRI data from 9 macaques were used to calculate fiber orientations using a ball-and-sticks model^65^. The fiber orientations were then registered to white matter masks derived from T2-weighted MRI. An edge-complete, but sparse tract-traced connectome^66^ was used to optimize the parameters of a probabilistic tractography algorithm^67^ applied to the masked fiber orientations. The whole cortex tractograms produced by this algorithm were then transformed to the F99 connectivity space^68^ and fiber density was used to estimate connection weights between the 82 regions of the regional map parcellation^69^. Finally, the resulting 82 by 82 weighted connectivity matrices were averaged across subjects, made symmetric and then constrained to the non-zero elements of a whole cortex tracer connectome^70^.

### Assigning Intrinsic Frequencies

Intrinsic activity within each of *n* cortical regions was drawn from the alpha band (8–12 Hz) and scaled by the strength of outgoing connections from that region^34,71^. For a region *i* with intrinsic frequency *w*_*i*_,

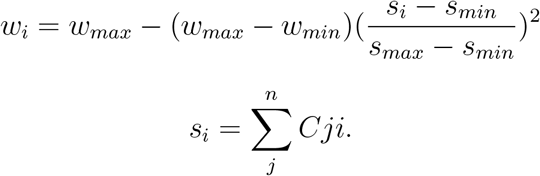

where *w*_*min*_ = 8 Hz, *w*_*max*_ = 12 Hz, *s*_*min*_, *s*_*max*_ are the lowest and highest region strengths and *C* is a structural connectivity matrix. For macaque simulations, *w*_*min*_ = 9 Hz and *w*_*max*_ = 11 Hz. This distribution of intrinsic frequencies is based on the hierarchy of timescales observed across the core-periphery axis^72,73^.

### Simulating Synchronization Dynamics

The human and macaque cortices were divided into regions based on the Schaefer parcellation^39^ (*n* = 1000) and the regional map parcellation^69^ (*n* = 82) respectively. Activity in each cortical region *i* was modeled as a combination of an intrinsic oscillation frequency (*w*_*i*_) and sinusoidal coupling to other regions, scaled by a structural connectivity matrix (*C*). Phase dynamics within each region (*θ*_*i*_) were simulated using the Kuramoto model^33^,

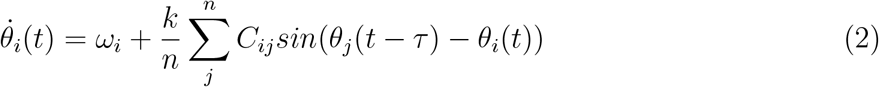

Activity was simulated with a time step of 1 millisecond and integrated using the Euler method. Synchronization between regions was quantified using the Kuramoto order parameter (*r*),

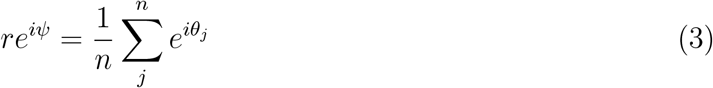

The scalar for global coupling (*k*) and the conduction delay (*τ*) were used to define the parameter space. A grid search was performed for *k* values from 0 to 8 and *τ* values from 1 to 15 in increments of 0.1. For each pair of parameters, 100 seconds of activity were simulated and a synchronization time series was quantified for the whole cortex. The frequency of slow coherence oscillation at each point in the parameter space was estimated by detecting local minima in the Kuramoto order parameter using peak detection and retaining only peaks with a minimum prominence of 0.4. Frequency was then computed as the number of detected minima divided by the total simulation duration. Synchronization was also calculated for the temporo-parietal and default-mode networks based on the functional networks of Yeo et al. ^40^ for humans and suitable analogs for macaques^43,44,45^. The scalar for global coupling (*k*) and the conduction delay (*τ*) were tuned to maximize variance in global *r* using a grid search. This is thought to index metastability, a feature of resting-state neural activity^46^. The search found optimal metastability at *k* = 2, *τ* = 4 for humans and *k* = 2.6,*τ* = 5 for macaques. The set of parameters maximizing metastability was used for any subsequent analysis. The same simulations were also performed on the AAL connectome, with connectivity matrices acquired from the public repository associated with Cabral et al. ^74^.

### Simulating Fluctuations in Stimulus Entrainment

Interactions between external input and network activity were modeled using a proxy region (*i*) with sinusoidal coupling to all regions within the temporo-parietal network and an external oscillatory input. Phase dynamics within the temporo-parietal network were isolated from whole cortex dynamics for both the human and macaque connectomes (see previous section). Activity in the proxy region was defined by the following equation,

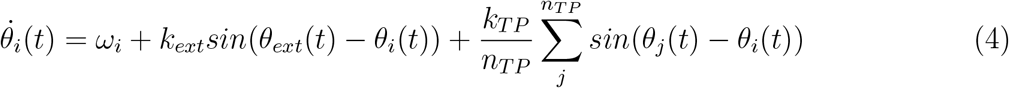

where *θ*_*i*_, *θ*_*ext*_ and *θ*_*j*_ are the phases of the stimulated region, the stimulus and a region in the temporo-parietal network respectively. *k*_*ext*_ and *k*_*TP*_ are scalars and *n*_*TP*_ is the number of regions in the temporo-parietal network. The intrinsic frequency of the proxy region (*w*_*i*_) is 5 Hz, the stimulus frequency is 3 Hz and the nodes within the temporo-parietal network have intrinsic frequencies between 8–12 Hz. The phase dynamics of this region were simulated in 1 millisecond increments and numerically integrated using the Euler method. A time-frequency decomposition was then performed using the continuous wavelet transform to study fluctuations in entrainment toward the stimulus.

### Simulating Cholinergic Gain

Cholinergic gain was simulated by multiplying the human and macaque structural connectivity matrices at all indices in either the left or right temporo-parietal network by a scalar. Whole cortex activity was then simulated (see Simulating Synchronization Dynamics). A temporoparietal network synchronization time series was generated with and without conduction delays. For the human connectome, *k* = 2, *τ* = 4 for the delayed simulation and *k* = 1.2, *τ* = 0 in the non-delayed condition. Macaque simulations used *k* = 2.6, *τ* = 5 and *k* = 2, *τ* = 0. The dominant frequency of the temporo-parietal network synchronization time series was estimated by detecting local minima in the Kuramoto order parameter using peak detection on the sign-inverted time series, retaining only peaks with a minimum prominence of 0.4. Frequency was then computed as the number of detected minima divided by the total simulation duration. This analysis was repeated for gain values between 1 and 5. For each gain value, the reported frequencies were averaged over the results of 5 runs with distinct initial phases.

### Simulating Synchronization Dynamics in Non-Modular Networks

Coherence dynamics were simulated under two non-modular structural topologies, using the previously described method (see Simulating Synchronization Dynamics). The first topology consisted of randomly permuted human and macaque structural connectivity matrices. The second topology comprised an all-to-all, homogeneously connected network (*n* = 1000) with natural frequencies drawn from a Lorentzian distribution centered at 10 Hz. Both structural topologies were used to generate coherence time series over a range of delay and global coupling values.

### Data and code availability

All code required to reproduce the simulations and analyses reported here, together with dependency specifications and instructions for reproducing the computational environment, is publicly available via the Open Science Framework project^75^ under the Creative Commons Attribution 4.0 International license. Access it with the following DOI: https://doi.org/10.17605/OSF.IO/24PMY.

The structural connectivity matrices used here were obtained from third-party public repositories and remain subject to the data-use terms of their original providers. The Schaefer-1000 structural connectivity matrix was obtained from the repository associated with Deco et al. ^47^. The macaque regional-map structural connectivity matrix was obtained from the public repository associated with Shen et al. ^42^. The AAL-90 structural connectivity matrix was obtained from the public repository associated with Cabral et al. ^74^. To avoid relicensing third-party data, the uploaded project files provide information for obtaining these matrices from their original repositories but do not redistribute them. Users are responsible for complying with the original data-use terms and for citing the original data sources. All data generated by our simulations, including processed outputs and figure data, are made available under the same project.

## CRediT authorship contribution statement

**Anagh Pathak**: Design, investigation, methodology and writing manuscript; **Rishabh Bapat**: Design, investigation, methodology and writing manuscript; **Arpan Banerjee**: Supervision, design, resources, reviewing and editing manuscript.

## Acknowledgments

The authors acknowledge the Computing Facility of the National Brain Research Centre, India for support with infrastructure and the Dementia Science Program awarded by the Department of Biotechnology, Government of India. Authors also acknowledge the [Neuroscience Gateway](https://ceur-ws.org/Vol-993/paper10.pdf)^76^ for assisting this work through the use of the Expanse supercomputer.

## References

[1] G. Buzsaki and A. Draguhn. Neuronal oscillations in cortical networks. science, 304(5679): 1926–1929, 2004.

[2] O. Jensen and L. L. Colgin. Cross-frequency coupling between neuronal oscillations. Trends in cognitive sciences, 11(7):267–269, 2007.

[3] N. J. Kopell, H. J. Gritton, M. A. Whittington, and M. A. Kramer. Beyond the connectome: The dynome. Neuron, 83(6):1319–1328, 2014.

[4] R. F. Betzel, M. Fukushima, Y. He, X.-N. Zuo, and O. Sporns. Dynamic fluctuations coincide with periods of high and low modularity in resting-state functional brain networks. NeuroImage, 127:287–297, February 2016. ISSN 1053-8119. doi: 10.1016/j.neuroimage.2015.12.001. URL https://www.sciencedirect.com/science/article/pii/S1053811915011143.

[5] D. Vidaurre, S. M. Smith, and M. W. Woolrich. Brain network dynamics are hierarchically organized in time. Proceedings of the National Academy of Sciences, 114(48):12827–12832, 2017.

[6] A. R. McIntosh. Contexts and catalysts: a resolution of the localization and integration of function in the brain. Neuroinformatics, 2(2):175–182, 2004. doi: 10.1385/NI:2:2:175.

[7] J. M. Shine, M. J. Aburn, M. Breakspear, and R. A. Poldrack. The modulation of neural gain facilitates a transition between functional segregation and integration in the brain. eLife, 7:e31130, 2018.

[8] T. Womelsdorf, T. A. Valiante, N. T. Sahin, K. J. Miller, and P. Tiesinga. Dynamic circuit motifs underlying rhythmic gain control, gating and integration. Nature Neuroscience, 17 (8):1031–1039, 2014.

[9] G. Deco, V. Jirsa, A. R. McIntosh, O. Sporns, and R. Kötter. Key role of coupling, delay, and noise in resting brain fluctuations. Proceedings of the National Academy of Sciences, 106(25):10302–10307, 2009.

[10] G. Tononi, O. Sporns, and G. M. Edelman. Measures of degeneracy and redundancy in biological networks. Proceedings of the National Academy of Sciences of the United States of America, 96(6):3257–3262, 1999. doi: 10.1073/pnas.96.6.3257. URL https://doi.org/10.1073/pnas.96.6.3257.

[11] K. Friston. Functional integration and inference in the brain. Prog Neurobiol, 68(2): 113–143, 2002. doi: 10.1016/s0301-0082(02)00076-x.

[12] O. Sporns. Structure and function of complex brain networks. Dialogues Clin Neurosci, 15(3), 2013.

[13] G. Deco, G. Tononi, M. Boly, and M. L. Kringelbach. Rethinking segregation and integration: Contributions of whole-brain modelling. Nature Reviews Neuroscience, 16(7): 430–439, 2015.

[14] S. S. Singh, A. Mukherjee, P. Raghunathan, D. Ray, and A. Banerjee. High segregation and diminished global integration in large-scale brain functional networks enhances the perceptual binding of cross-modal stimuli. Cerebral Cortex, 34(8):bhae323, 2024. doi: 10.1093/cercor/bhae323.

[15] J. R. Cohen and M. D’Esposito. The segregation and integration of distinct brain networks and their relationship to cognition. Journal of Neuroscience, 36(48):12083–12094, 2016.

[16] J. M. Shine. Neuromodulatory influences on integration and segregation in the brain. Trends in cognitive sciences, 23(7):572–583, 2019.

[17] S. L. Bressler, R. Coppola, and R. Nakamura. Episodic multiregional cortical coherence at multiple frequencies during visual task performance. Nature, 366(6451):153–156, 1993. doi: 10.1038/366153a0.

[18] J. M. Shine, P. G. Bissett, P. T. Bell, O. Koyejo, J. H. Balsters, K. J. Gorgolewski, C. A. Moodie, and R. A. Poldrack. The dynamics of functional brain networks: Integrated network states during cognitive task performance. Neuron, 92(2):544–554, October 2016. ISSN 0896-6273. doi: 10.1016/j.neuron.2016.09.018. URL https://www.sciencedirect.com/science/article/pii/S0896627316305773.

[19] R. Wang, M. Liu, X. Cheng, Y. Wu, A. Hildebrandt, and C. Zhou. Segregation, integration, and balance of large-scale resting brain networks configure different cognitive abilities. Proceedings of the National Academy of Sciences, 118(23):e2022288118, 2021.

[20] A. Zalesky, A. Fornito, L. Cocchi, L. L. Gollo, and M. Breakspear. Time-resolved resting-state brain networks. Proceedings of the National Academy of Sciences, 111(28):10341– 10346, June 2014. ISSN 1091-6490. doi: 10.1073/pnas.1400181111.

[21] R. Liégeois, E. Ziegler, C. Phillips, P. Geurts, F. Gómez, M. A. Bahri, B. T. T. Yeo, A. Soddu, A. Vanhaudenhuyse, S. Laureys, and R. Sepulchre. Cerebral functional connectivity periodically (de)synchronizes with anatomical constraints. Brain Structure and Function, 221(6):2985–2997, 2016. ISSN 1863-2653. doi: 10.1007/s00429-015-1083-y.

[22] Z.-Q. Gong and X.-N. Zuo. Dark brain energy: Toward an integrative model of spontaneous slow oscillations. Physics of Life Reviews, 52:278–297, 2025.

[23] J. C. Pang. Fundamental constraints shaping intrinsic brain activity: Comment on “Dark brain energy: Toward an integrative model of spontaneous slow oscillations” by Zhu-Qing Gong and Xi-Nian Zuo. Physics of Life Reviews, 2025.

[24] M. W. Cole, D. S. Bassett, J. D. Power, T. S. Braver, and S. E. Petersen. Intrinsic and task-evoked network architectures of the human brain. Neuron, 83(1):238–251, 2014. doi: 10.1016/j.neuron.2014.05.014.

[25] C. J. Honey, E. L. Newman, and A. C. Schapiro. Switching between internal and external modes: A multiscale learning principle. Network Neuroscience, 1(4):339–356, 2017. ISSN 2472-1751. doi: 10.1162/netna00024.

[26] A. C. Nobre and D. Gresch. How the brain shifts between external and internal attention. Neuron, 113(15):2382–2398, 2025.

[27] P. Lakatos, A. Barczak, S. A. Neymotin, T. McGinnis, D. Ross, D. C. Javitt, and M. N. O’Connell. Global dynamics of selective attention and its lapses in primary auditory cortex. Nature Neuroscience, 19(12):1707–1717, 2016. ISSN 1097-6256. doi: 10.1038/nn.4386.

[28] F. H. Kasten, Q. Busson, and B. Zoefel. Opposing neural processing modes alternate rhythmically during sustained auditory attention. Communications Biology, 7(1):1125, September 2024. ISSN 2399-3642. doi: 10.1038/s42003-024-06834-x. URL https://www.nature.com/articles/s42003-024-06834-x.

[29] R. G. OConnell, P. M. Dockree, I. H. Robertson, M. A. Bellgrove, J. J. Foxe, and S. P. Kelly. Uncovering the Neural Signature of Lapsing Attention: Electrophysiological Signals Predict Errors up to 20 s before They Occur. Journal of Neuroscience, 29(26):8604–8611, July 2009. ISSN 0270-6474, 1529-2401. doi: 10.1523/JNEUROSCI.5967-08.2009. URL https://www.jneurosci.org/content/29/26/8604.

[30] A. Vanhaudenhuyse, A. Demertzi, M. Schabus, Q. Noirhomme, S. Bredart, M. Boly, C. Phillips, A. Soddu, A. Luxen, G. Moonen, and S. Laureys. Two distinct neuronal networks mediate the awareness of environment and of self. Journal of Cognitive Neuroscience, 23(3):570–578, March 2011. ISSN 1530-8898. doi: 10.1162/jocn.2010.21488.

[31] M. Bastian and J. Sackur. Mind wandering at the fingertips: Automatic parsing of subjective states based on response time variability. Frontiers in Psychology, 4, September 2013. ISSN 1664-1078. doi: 10.3389/fpsyg.2013.00573. URL https://www.frontiersin.org/journals/psychology/articles/10.3389/fpsyg.2013.00573/full.

[32] D. Marr. Vision: A Computational Investigation into the Human Representation and Processing of Visual Information. MIT Press, Cambridge, MA, 2010.

[33] Y. Kuramoto. Chemical Oscillations, Waves, and Turbulence. Springer Berlin Heidelberg, 1984. ISBN 978-3-642-69689-3. doi: 10.1007/978-3-642-69689-3.

[34] A. Pathak, V. Sharma, D. Roy, and A. Banerjee. Biophysical mechanism underlying compensatory preservation of neural synchrony over the adult lifespan. Communications Biology, 5(1):567, 2022.

[35] P. Ritter, M. Schirner, A. R. McIntosh, and V. K. Jirsa. The virtual brain integrates computational modeling and multimodal neuroimaging. Brain Connectivity, 3(2):121–145, 2013. doi: 10.1089/brain.2012.0120.

[36] A. Pathak, D. Roy, and A. Banerjee. Whole-brain network models: From physics to bedside. Frontiers in Computational Neuroscience, 16:866517, 2022.

[37] R. Bapat, A. Pathak, and A. Banerjee. Metastability indexes global changes in the dynamic working point of the brain following brain stimulation. Frontiers in Neurorobotics, 18: 1336438, 2024.

[38] H. Haken. Synergetics: An Introduction. Springer Verlag, 1983.

[39] A. Schaefer, R. Kong, E. M. Gordon, T. O. Laumann, X.-N. Zuo, A. J. Holmes, S. B. Eickhoff, and B. T. T. Yeo. Local-global parcellation of the human cerebral cortex from intrinsic functional connectivity MRI. Cerebral Cortex, 28(9):3095–3114, July 2017. ISSN 1460-2199. doi: 10.1093/cercor/bhx179.

[40] B. T. T. Yeo, F. M. Krienen, J. Sepulcre, M. R. Sabuncu, D. Lashkari, M. Hollinshead, J. L. Roffman, J. W. Smoller, L. Zöllei, J. R. Polimeni, B. Fischl, H. Liu, and R. L. Buckner. The organization of the human cerebral cortex estimated by intrinsic functional connectivity. Journal of Neurophysiology, 106(3):1125–1165, September 2011. ISSN 0022-3077. doi: 10.1152/jn.00338.2011. URL https://www.ncbi.nlm.nih.gov/pmc/articles/PMC3174820/.

[41] E. T. Rolls, C.-C. Huang, C.-P. Lin, J. Feng, and M. Joliot. Automated anatomical labelling atlas 3. NeuroImage, 206:116189, February 2020. ISSN 1053-8119. doi: 10.1016/j.neuroimage.2019.116189. URL https://www.sciencedirect.com/science/article/pii/S1053811919307803.

[42] K. Shen, G. Bezgin, M. Schirner, P. Ritter, S. Everling, and A. R. McIntosh. A macaque connectome for large-scale network simulations in TheVirtualBrain. Scientific Data, 6(1): 123, July 2019. ISSN 2052-4463. doi: 10.1038/s41597-019-0129-z. URL https://www.nature.com/articles/s41597-019-0129-z.

[43] E. Borra and G. Luppino. Functional anatomy of the macaque temporo-parieto-frontal connectivity. Cortex; a journal devoted to the study of the nervous system and behavior, 97:306–326, December 2017. ISSN 0010-9452. doi: 10.1016/j.cortex.2016.12.007. URL https://www.sciencedirect.com/science/article/pii/S0010945216303513.

[44] A. Touroutoglou, E. Bliss-Moreau, J. Zhang, D. Mantini, W. Vanduffel, B. C. Dickerson, and L. F. Barrett. A ventral salience network in the macaque brain. NeuroImage, 132: 190–197, May 2016. ISSN 1053-8119. doi: 10.1016/j.neuroimage.2016.02.029. URL https://www.sciencedirect.com/science/article/pii/S1053811916001361.

[45] D. Mantini, A. Gerits, K. Nelissen, J.-B. Durand, O. Joly, L. Simone, H. Sawamura, C. Wardak, G. A. Orban, R. L. Buckner, et al. Default mode of brain function in monkeys. Journal of Neuroscience, 31(36):12954–12962, 2011.

[46] E. Tognoli and J. A. S. Kelso. The metastable brain. Neuron, 81(1):35–48, 2014.

[47] G. Deco, M. L. Kringelbach, V. K. Jirsa, and P. Ritter. The dynamics of resting fluctuations in the brain: Metastability and its dynamical cortical core. Scientific Reports, 7(1):3095, 2017.

[48] E. Ott and T. M. Antonsen. Low dimensional behavior of large systems of globally coupled oscillators. Chaos: An Interdisciplinary Journal of Nonlinear Science, 18(3):037113, 2008.

[49] P. Villegas, P. Moretti, and M. A. Muñoz. Frustrated hierarchical synchronization and emergent complexity in the human connectome network. Scientific Reports, 4(1), August 2014. ISSN 2045-2322. doi: 10.1038/srep05990.

[50] U. Alon. Network motifs: theory and experimental approaches. Nat Rev Genet, 8(6): 450–461, 2007. doi: 10.1038/nrg2102.

[51] J. C. Pang, K. M. Aquino, M. Oldehinkel, P. A. Robinson, B. D. Fulcher, M. Breakspear, and A. Fornito. Geometric constraints on human brain function. Nature, 618(7965):566–574, 2023.

[52] 1 F. Milisav, A. I. Luppi, L. E. Suárez, G. Lajoie, and B. Misic. Neuromorphic hierarchical modular reservoirs. Nature Communications, 2026.

[53] M. Glukhova, A. Tlaie, R. Taylor, P.-A. Ferracci, K. Shapcott, B. Mert, O. Arne, A. Ciuparu, R. C. Muresan, M. N. Havenith, and M. L. Schölvinck. Sharing the spotlight: Uncovering common attentional dynamics across species. PLOS Computational Biology, 22(4): e1014191, April 2026. ISSN 1553-7358. doi: 10.1371/journal.pcbi.1014191. URL https://journals.plos.org/ploscompbiol/article?id=10.1371/journal.pcbi.1014191.

[54] G. Vallortigara, L. J. Rogers, and A. Bisazza. Possible evolutionary origins of cognitive brain lateralization. Brain Research Reviews, 30(2):164–175, 1999.

[55] M. C. Corballis. Left brain, right brain: Facts and fantasies. PLOS Biology, 12(1): e1001767, 2014.

[56] M. Corbetta and G. L. Shulman. Spatial neglect and attention networks. Annual review of neuroscience, 34(1):569–599, 2011.

[57] M. Thiebaut de Schotten and C. F. Beckmann. Asymmetry of brain structure and function: 40 years after Sperry’s Nobel Prize. Brain Structure and Function, 227(2):421–424, 2022.

[58] M. L. Concha, I. H. Bianco, and S. W. Wilson. Encoding asymmetry within neural circuits. Nature Reviews Neuroscience, 13(12):832–843, 2012.

[59] N. He, L. Palaniyappan, Z. Linli, and S. Guo. Abnormal hemispheric asymmetry of both brain function and structure in attention deficit/hyperactivity disorder: A meta-analysis of individual participant data. Brain Imaging and Behavior, 16(1):54–68, 2022.

[60] D. Li, T. Li, Y. Niu, J. Xiang, R. Cao, B. Liu, H. Zhang, and B. Wang. Reduced hemispheric asymmetry of brain anatomical networks in attention deficit hyperactivity disorder. Brain Imaging and Behavior, 13(3):669–684, 2019.

[61] D. C. Van Essen, S. M. Smith, D. M. Barch, T. E. J. Behrens, E. Yacoub, K. Ugurbil, Wu-Minn HCP Consortium, et al. The WU-Minn human connectome project: An overview. NeuroImage, 80:62–79, 2013.

[62] F.-C. Yeh, V. J. Wedeen, and W.-Y. I. Tseng. Generalized q-sampling imaging. IEEE transactions on medical imaging, 29(9):1626–1635, 2010.

[63] F.-C. Yeh, T. D. Verstynen, Y. Wang, J. C. Fernández-Miranda, and W.-Y. I. Tseng. Deterministic diffusion fiber tracking improved by quantitative anisotropy. PloS one, 8 (11):e80713, 2013.

[64] J. Mazziotta, A. Toga, A. Evans, P. Fox, J. Lancaster, K. Zilles, R. Woods, T. Paus, G. Simpson, B. Pike, et al. A four-dimensional probabilistic atlas of the human brain. Journal of the American Medical Informatics Association, 8(5):401–430, 2001.

[65] S. Jbabdi, S. N. Sotiropoulos, A. M. Savio, M. Graña, and T. E. J. Behrens. Model-based analysis of multishell diffusion MR data for tractography: How to get over fitting problems. Magnetic resonance in medicine, 68(6):1846–1855, 2012.

[66] N. T. Markov, M. M. Ercsey-Ravasz, A. R. Ribeiro Gomes, C. Lamy, L. Magrou, J. Vezoli, P. Misery, A. Falchier, R. Quilodran, M. A. Gariel, J. Sallet, R. Gamanut, C. Huissoud, S. Clavagnier, P. Giroud, D. Sappey-Marinier, P. Barone, C. Dehay, Z. Toroczkai, aK. Knoblauch, D. C. Van Essen, and H. Kennedy. A Weighted and Directed Interareal Connectivity Matrix for Macaque Cerebral Cortex. Cerebral Cortex, 24(1):17–36, January 2014. ISSN 1047-3211. doi: 10.1093/cercor/bhs270. URL https://doi.org/10.1093/cercor/bhs270.

[67] T. E. J. Behrens, H. J. Berg, S. Jbabdi, M. F. S. Rushworth, and M. W. Woolrich. Probabilistic diffusion tractography with multiple fibre orientations: What can we gain? NeuroImage, 34(1):144–155, 2007.

[68] D. C. Van Essen and D. L. Dierker. Surface-based and probabilistic atlases of primate cerebral cortex. Neuron, 56(2):209–225, 2007.

[69] R. Kötter and E. Wanke. Mapping brains without coordinates. Philosophical Transactions of the Royal Society B: Biological Sciences, 360(1456):751–766, April 2005. doi: 10.1098/rstb.2005.1625. URL https://royalsocietypublishing.org/doi/10.1098/rstb.2005.1625.

[70] K. Shen, G. Bezgin, R. M. Hutchison, J. S. Gati, R. S. Menon, S. Everling, and A. R. McIntosh. Information processing architecture of functionally defined clusters in the macaque cortex. Journal of Neuroscience, 32(48):17465–17476, 2012.

[71] L. Cocchi, M. V. Sale, L. L. Gollo, P. T. Bell, V. T. Nguyen, A. Zalesky, M. Breakspear, and J. B. Mattingley. A hierarchy of timescales explains distinct effects of local inhibition of primary visual cortex and frontal eye fields. eLife, 5:e15252, 2016.

[72] J. D. Murray, A. Bernacchia, D. J. Freedman, R. Romo, J. D. Wallis, X. Cai, C. Padoa-Schioppa, T. Pasternak, H. Seo, D. Lee, et al. A hierarchy of intrinsic timescales across primate cortex. Nature Neuroscience, 17(12):1661–1663, 2014.

[73] L. L. Gollo, A. Zalesky, R. M. Hutchison, M. Van Den Heuvel, and M. Breakspear. Dwelling quietly in the rich club: Brain network determinants of slow cortical fluctuations. Philosophical Transactions of the Royal Society B: Biological Sciences, 370(1668):20140165, 2015.

[74] J. Cabral, H. Luckhoo, M. Woolrich, M. Joensson, H. Mohseni, A. Baker, M. L. Kringelbach, and G. Deco. Exploring mechanisms of spontaneous functional connectivity in MEG: How delayed network interactions lead to structured amplitude envelopes of bandpass filtered oscillations. NeuroImage, 90:423–435, April 2014. ISSN 1095-9572. doi: 10.1016/j.neuroimage.2013.11.047.

[75] E. D. Foster and A. Deardorff. Open science framework (osf). Journal of the Medical Library Association: JMLA, 105(2):203, 2017.

[76] S. Sivagnanam, A. Majumdar, K. Yoshimoto, V. Astakhov, A. Bandrowski, M. Martone, and N. Carnevale. Introducing the neuroscience gateway. In IWSG, 2013. URL https://www.semanticscholar.org/paper/9e0b5dc3dc5c389526722b1ee856afdcdd7a6468.

